# Activation by statins unveils two putative agonist binding sites in the pore domain of TRPA1

**DOI:** 10.64898/2026.05.08.723702

**Authors:** Justyna B. Startek, Alina Milici, Katharina Held, Ariel Talavera, Karel Talavera

## Abstract

TRPA1 is a non-selective cation channel that plays a crucial role in several pain and inflammatory conditions. Agents reducing membrane cholesterol decrease TRPA1 activation, but it remains unclear how cholesterol-lowering medications affect TRPA1 function. Given that TRPA1 is activated by a wide variety of chemicals, we explored whether statins have acute effects on this channel. We found that five commonly used statins activate human and mouse TRPA1 in a reversible and concentration-dependent manner. The effective concentrations were above the micromolar range, in the order: simvastatin ≈ lovastatin < fluvastatin < atorvastatin < pravastatin. Statin-induced activation was not correlated to changes in membrane order, nor mediated by N-terminal cysteine residues contributing to electrophilic compound agonism. Molecular docking calculations and the functional characterization of single-point mutants revealed two separate putative binding sites, one situated close to the kink of transmembrane segment 5 (TM5) and the other at the interface between TM4 and TM5. The mTRPA1 inhibitor A-967079 largely abrogated the response to the electrophilic agonist allyl isothiocyanate, but had weaker and varied effects across different statins and menthol. Mutation T877L strongly altered the effect of A-967079, also in an agonist-dependent manner, suggesting competitive binding between this antagonist and the non-electrophilic agonists. The identification of two distinct agonist binding sites may help explaining how TRPA1 is able to respond to a large variety of non-electrophilic compounds, while the finding of competitive interactions at one of these sites may help guide the development of agonist-specific antagonists of therapeutic relevance.

## Introduction

Sensory neurons express various receptors that detect chemical irritants and harmful thermal and mechanical stimuli, many of which belong to the Transient Receptor Potential (TRP) cation channel superfamily. TRP channel activation in nociceptors results in cation influx, leading to electrical excitation and pain sensation, and to the release of neuropeptides that trigger local inflammatory responses [1]. One member of the TRP superfamily, the ankyrin-rich TRPA1, is arguably the most versatile polymodal sensor, being activated by an impressive variety of noxious chemicals, and by thermal and mechanical stimuli [2, 3].

Previously, we showed that TRPA1 gating and expression in lipid rafts is modulated by cholesterol, which can interact directly with the channel through cholesterol recognition amino acid consensus (CRAC) motifs located in the transmembrane segments TM2 and TM4 [4]. Cholesterol depletion induced with methyl-β-cyclodextrin (MCD) or sphingomyelinase (SMase) decreases TRPA1 responses to chemical agonists such as AITC and thymol, to bacterial lipopolysaccharides, and to cold [4, 5]. Moreover, cholesterol may have played a permissive role in the responses of reconstituted TRPA1 channels to chemical, thermal, and mechanical stimuli in artificial lipid bilayers [6–9]. These findings raised the question of whether cholesterol-reducing drugs that are used in the clinic may modulate TRPA1 function.

Statins are the most relevant class of cholesterol-depleting drugs. They are HMG-CoA reductase inhibitors commonly used for long-term treatment of hypercholesterolemia and mixed hyperlipidemia, effectively reducing the risk of heart attacks [10–13]. Of note, statins affect cholesterol synthesis virtually in all cells, where they can produce increased bilayer heterogeneity and reorganization of discrete membrane domains [14], such as lipid rafts where TRPA1 is expressed [4].

The effects of statins on TRPA1 remain largely understudied, with one report showing that simvastatin did not affect Ca^2+^ influx in HEK293 cells transfected with human TRPA1 [15], and another one showing activation by simvastatin, and to a lesser extent by atorvastatin and rosuvastatin [16]. Motivated by these contradictory reports in the literature on hTRPA1, in this study we (re-)evaluated the effects of five statins on human and mouse TRPA1. We found concentration-dependent activation of human and mouse TRPA1 by simvastatin, lovastatin, fluvastatin, atorvastatin and pravastatin. Next, we investigated the mechanism of action in mTRPA1, known to have a more complex pharmacology than that of hTRPA1 [2]. Our results are not consistent with activation of mTRPA1 by mechanical perturbations in the plasma membrane, nor by covalent modification of N-terminal cysteine residues, but by direct interactions of statins with two distinct binding sites formed by amino acid residues of the 4^th^, 5^th^ and 6^th^ transmembrane segments. Furthermore, we show evidence that one of these putative sites can accommodate agonists and inhibitors simultaneously.

## Results

### Statins activate human and mouse TRPA1 channels

To evaluate the effects of statins on human TRPA1 we performed intracellular Ca^2+^ imaging in HEK293 cells transfected with this channel. We tested five statins commonly prescribed to reduce patient cholesterol levels: simvastatin, lovastatin, fluvastatin, atorvastatin and pravastatin. We chose statins characterized by different lipophilicity, as determined by the calculated octanol-water distribution coefficient (logP_ow_) (Table 1). Lipophilicity is a proxy for the ability of the compounds to insert in the plasma membrane, to modulate its fluidity, and to access binding sites in the channel. All statins tested induced an increase in intracellular Ca^2+^ (Figure 1a-e), in a concentration-dependent manner, with higher concentrations required for the least lipophilic statins (Figure 1f). We were unable to estimate the EC_50_ for pravastatin as it induced substantial changes in cell morphology and detachment from the coverslip at concentrations above 1000 µM. All cells responding to the statins were also stimulated by the TRPA1 agonist allyl-isothiocyanate (AITC, 100 µM), and non-transfected HEK293 cells did not respond to any of the statins (Figure 1a-e).

**Figure 1.**
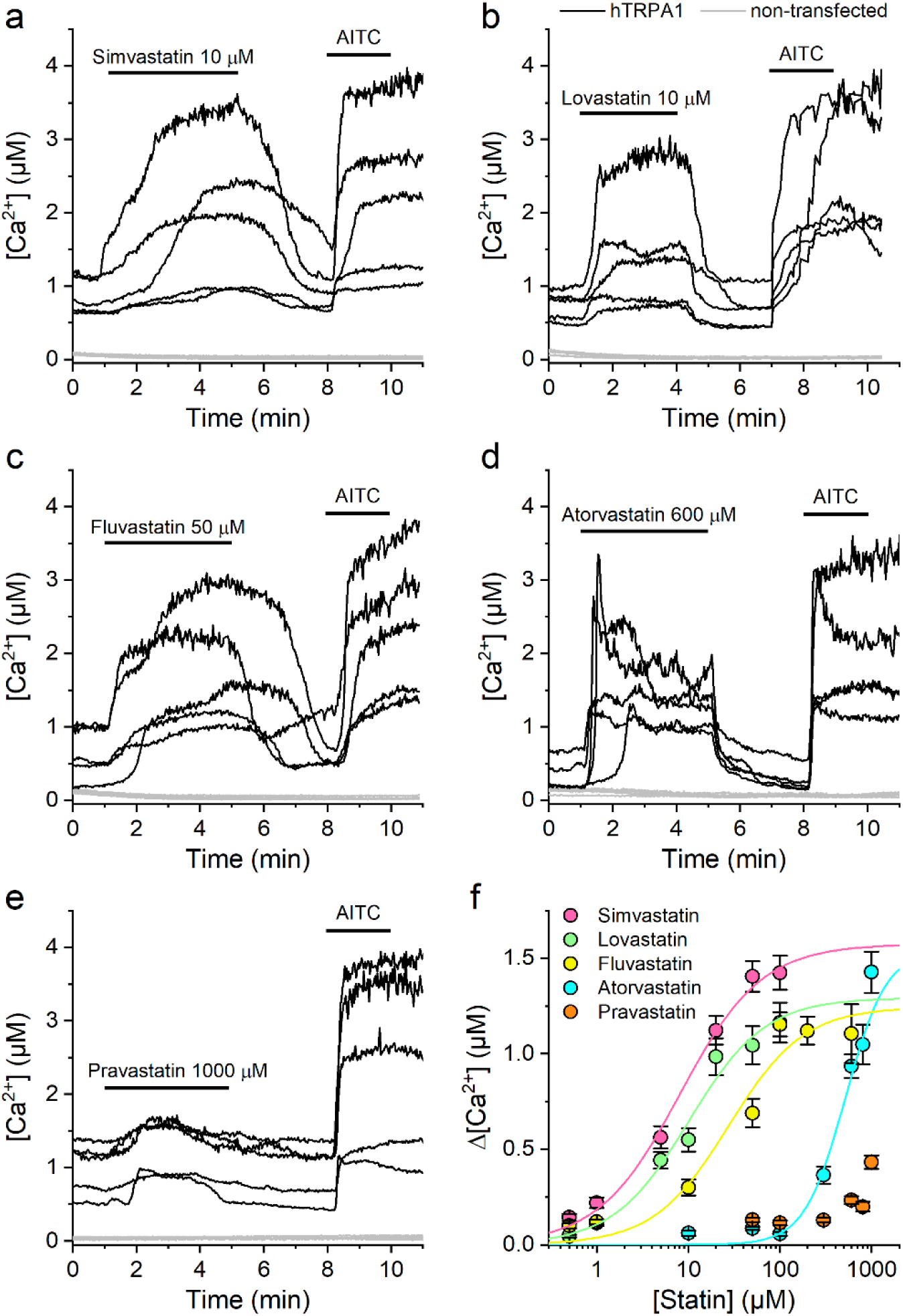
Statins activate hTRPA1 in a concentration-dependent manner. (a-e) Representative traces of [Ca^2+^] recorded in HEK-hTRPA1 or non-transfected HEK293 cells showing the effects of statins applied at concentrations near EC_50_ for hTRPA1 activation: simvastatin (10 µM, a), lovastatin (10 µM, b), fluvastatin (50 µM, c), atorvastatin (600 µM, d) and pravastatin (1000 µM, e). (f) Dose-response curves were derived from [Ca^2+^] responses (peak fluorescence baseline corrected) of hTRPA1 to statins, from which compounds EC_50_ values were determined. EC_50_ in µM: 7.9 ± 1, simvastatin; 10.1 ± 1, lovastatin; 28.2 ± 6, fluvastatin; 483.7 ± 37, atorvastatin; >1000, pravastatin. Data from n>100 cells are shown as mean ± s.e.m. The solid lines represent the fit of the data with Hill functions, yielding EC_50_ value ± fitting error.

**Table 1.**
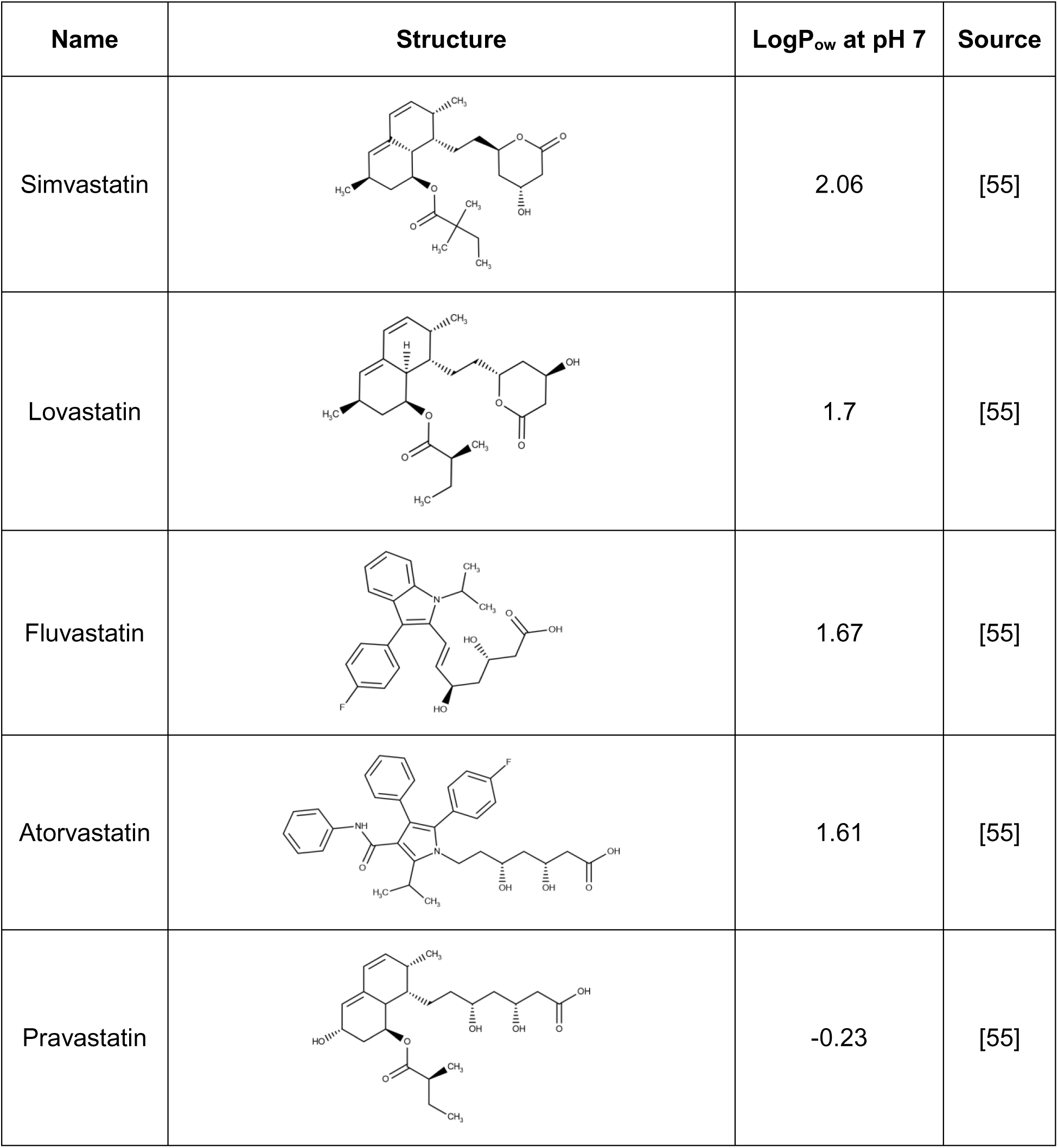
List of reagents used in the study, structure, and experimental partition coefficient (logP_ow_). The purity is above 98% for all compounds.

| Name | Structure | $\log P_{ow}$ at pH 7 | Source |
| --- | --- | --- | --- |
| Simvastatin |  | 2.06 | [55] |
| Lovastatin |  | 1.7 | [55] |
| Fluvastatin |  | 1.67 | [55] |
| Atorvastatin |  | 1.61 | [55] |
| Pravastatin |  | -0.23 | [55] |

The effects of multiple TRPA1 agonists and antagonists vary across orthologs of different species [2]. Thus, we tested whether activation of TRPA1 channels by statins is conserved between the human and mouse orthologs. We first performed intracellular Ca^2+^ imaging in Chinese Hamster Ovary (CHO) cells stably transfected with this channel (CHO-mTRPA1). As for hTRPA1, all statins increased the intracellular Ca^2+^ concentration (Figure 2a-e), but higher concentrations were required for the least lipophilic ones (Figure 2f). According to the EC_50_ values estimated from the fit of the experimental data by a Hill function, the potencies followed the order simvastatin ≈ lovastatin > fluvastatin > atorvastatin > pravastatin (Figure 2f). AITC (100 µM) stimulated all cells responding to the statins. In contrast, neither non-transfected CHO cells nor CHO-mTRPA1 treated with the mTRPA1 inhibitor HC-030031 (100 µM) (Figure 2a-e and Figure 2-figure supplement 1) responded to statins.

**Figure 2.**
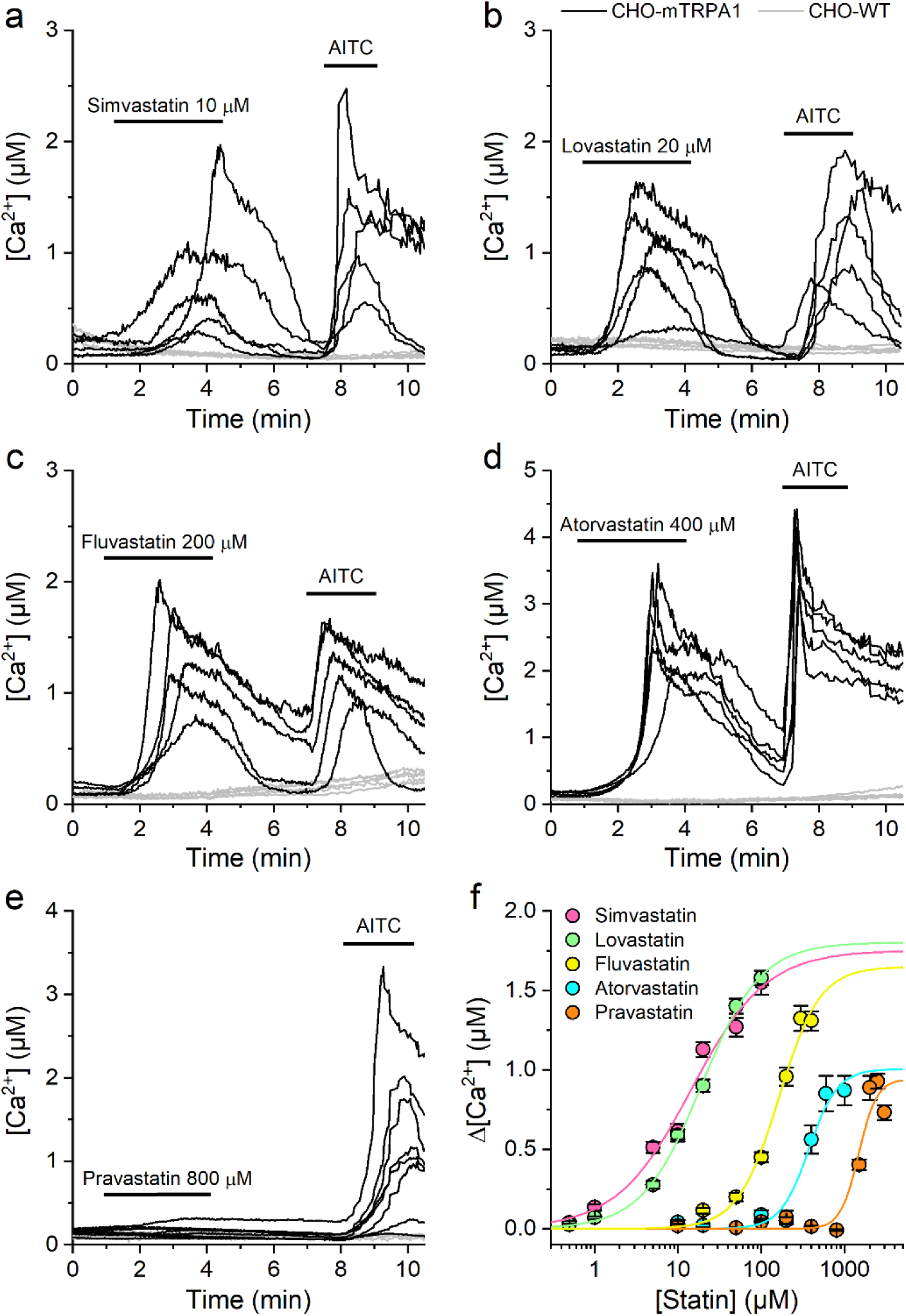
Statins activate mTRPA1 in a concentration-dependent manner. (a-e) Representative traces of [Ca^2+^] recorded in CHO-mTRPA1 (black traces) or non-transfected CHO cells (light gray traces) showing the effects of simvastatin (10 µM, a), lovastatin (20 µM, b), fluvastatin (120 µM, c), atorvastatin (400 µM, d), and pravastatin (800 µM, e). (f) Concentration-response curves derived from [Ca^2+^] responses (peak fluorescence baseline corrected) of mTRPA1 to statins, from which compounds EC_50_ values were determined. EC_50_ in µM: 14.4 ± 5, simvastatin; 19.1 ± 2, lovastatin; 162.7 ± 39, fluvastatin; 375.8 ± 53, atorvastatin; 1470 ± 181, pravastatin. Data from n > 80 cells shown as mean ± s.e.m.

We further tested the effects of statins on mTRPA1 by performing whole-cell patch-clamp experiments in CHO-mTRPA1 cells. Extracellular application of 10 µM simvastatin, 20 µM lovastatin, 120 µM fluvastatin, 360 µM atorvastatin, and 800 µM pravastatin reversibly increased inward and outward currents (Figure 3), preserving the double rectification pattern and inactivation at very positive potentials, characteristic of TRPA1 currents (Figure 3a-c, insets). After a period of washout of the statins, subsequent applications of AITC (100 µM) produced stronger current increase.

**Figure 3.**
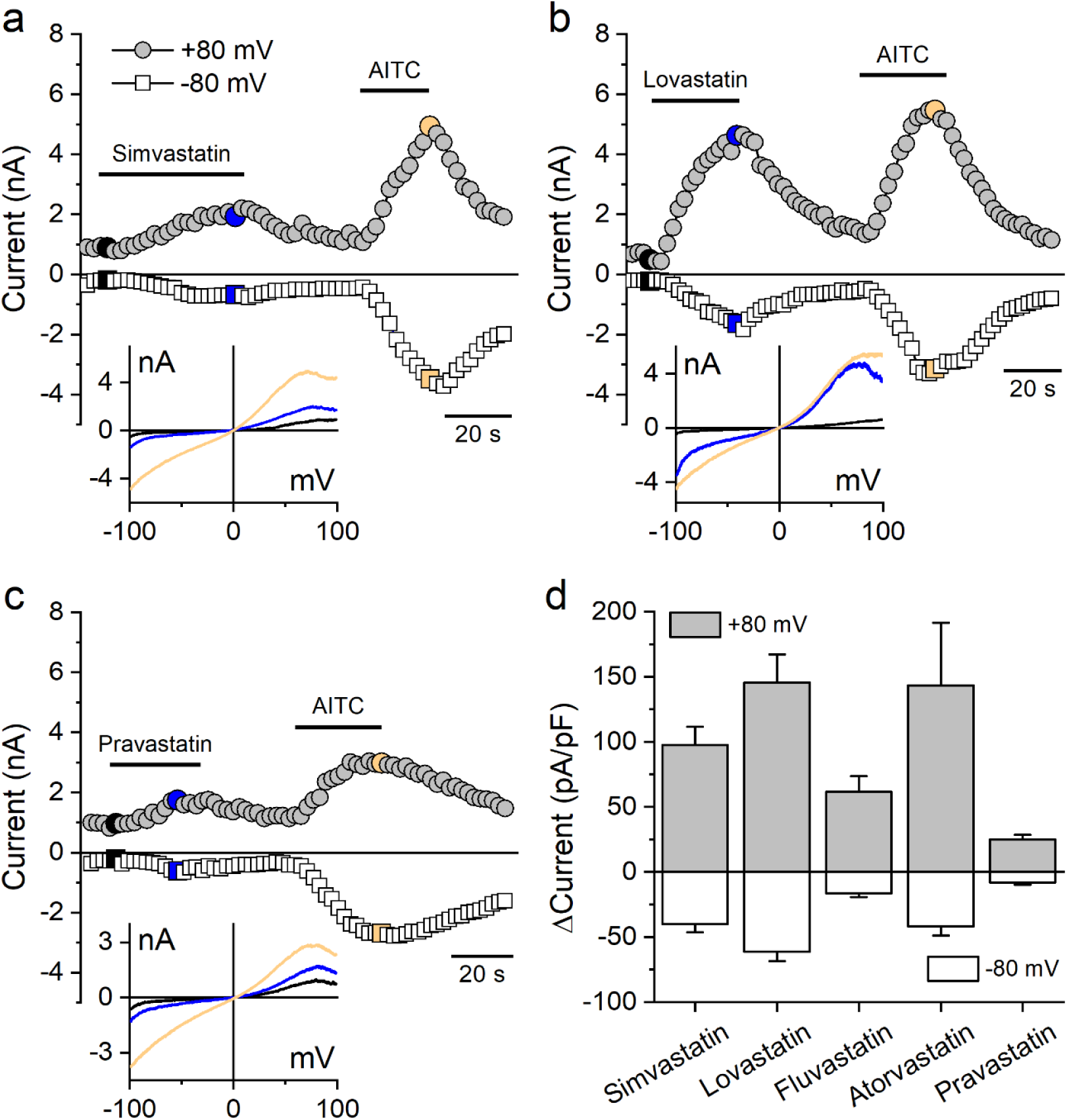
Statins stimulate mTRPA1 currents. (a-c) Time course of whole-cell patch-clamp recordings at holding potentials of +80 mV (gray circles) and −80 mV (white circles) of mTRPA1 expressed in CHO cells during application of statins, simvastatin (10 μM), lovastatin (20 μM), and pravastatin (800 μM). The insets in the bottom left corner display the current-voltage relationship (I–V) at the time points indicated in the main panels. (d) Statistics of mTRPA1 current increases at ±80 mV induced by different statins. Data are shown as mean ± s.e.m for n = 6-7 for each treatment.

Next, we determined the effects of statins in mouse DRGs, and the possible contribution of TRPA1 in this native expression system. Using Ca^2+^ imaging, we found that 10 µM simvastatin, 20 µM lovastatin, 120 µM fluvastatin, 360 µM atorvastatin, and 800 µM pravastatin stimulated 12.1% (93/769), 14.2% (87/612), 58.4% (506/866), 11.6% (66/571), and 2.3% (16/687) of neurons, respectively. Notably, a significant fraction of DRG neurons that responded to statins also responded to TRPA1 agonists 30 µM AITC (40%, 71%, 47% and 77% respectively to simva-, lova-, fluva- and atorvastatin) and 300 μM cinnamaldehyde (CA; 42%, 63%, 46% and 85% respectively to simva-, lova-, fluva- and atorvastatin) and TRPV1 activator 1 µM capsaicin (47%, 91%, 80% and 86% respectively to simva-, lova-, fluva- and atorvastatin) (Figure 4a-e). The selective TRPA1 antagonist HC-030031 largely prevented statin-induced Ca^2+^ responses (p < 0.0001 for all statins tested), and its washout in the presence of the statins led to strong responses (Figure 4-figure supplement 1).

**Figure 4.**
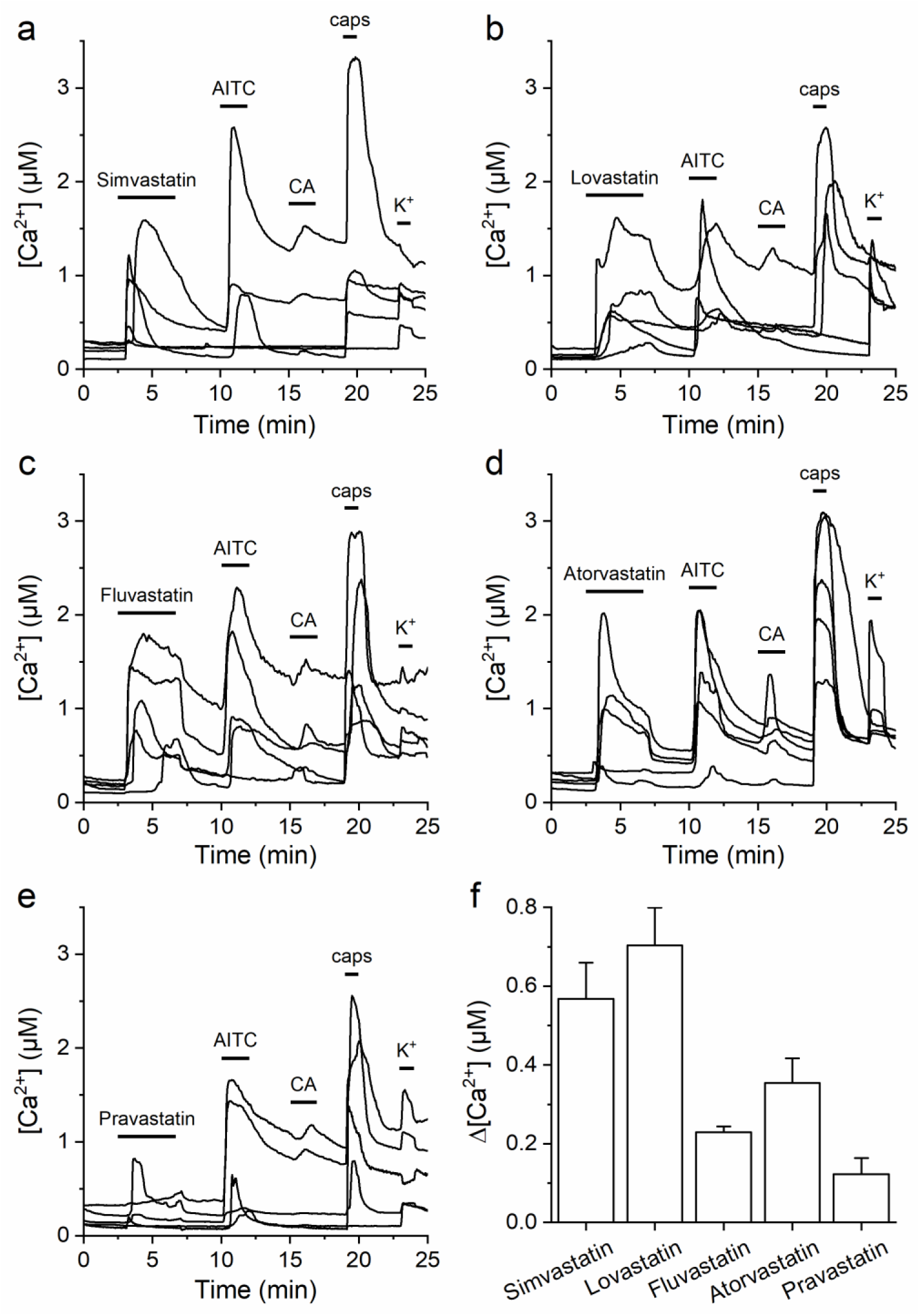
Statins activate TRP channels in mouse sensory neurons. (a-e) Representative traces of [Ca^2+^] responses of mouse DRG neurons to (a) 10 µM simvastatin; (b) 20 µM lovastatin; (c) 120 µM fluvastatin; (d) 360 µM atorvastatin; (e) 800 µM pravastatin and 30 µM AITC; 300 µM CA; 1 µM caps and high K^+^ (50 mM). (f) Average [Ca^2+^] amplitude change evoked by statins. Data are shown as mean ± s.e.m for n = 35, 59, 318, 49, 13 cells that were positive to simvastatin, lovastatin, fluvastatin, atorvastatin, and pravastatin, respectively, as well as to AITC and K^+^.

### TRPA1 activation does not correlate with plasma membrane fluidization induced by statins

Several studies have demonstrated that TRPA1 is sensitive to its membrane environment and that various non-electrophilic agonists TRPA1 reduce membrane fluidity [4, 5, 22–24]. Given that the agonist action of statins could be related to their lipophilicity, we addressed the possibility that they induce plasma membrane perturbations. We used Laurdan to test whether the partition of statins in the plasma membrane of CHO-mTRPA1 cells induces changes in the bilayer lipid order. This dye displays a red-shifted emission when in contact with water molecules that penetrate the membrane upon an increase in local lipid disorder [25]. These alterations can be quantified by changes in the Generalized Polarisation parameter determined at 340 nm (ΔexGP^340^) [22, 24–26]. As anticipated, benzyl alcohol (BA), a known bilayer disordering agent [27], strongly decreased exGP^340^ (Figure 5a). Treatment with simvastatin, lovastatin, fluvastatin, or atorvastatin (100 µM) for 5 min decreased exGP^340^, indicating membrane fluidization (Figure 5a). We further evaluated the effects of statins on the plasma membrane using diphenylhexatriene (DPH), a dye whose fluorescence anisotropy is reduced upon increases in membrane fluidity. BA application decreased anisotropy, as expected from its known fluidizing action (Figure 5b). Treatment with statins for 5 min induced a decrease in anisotropy, consistent with membrane fluidization (Figure 5b). We next enquired on whether the agonist action of the statins on TRPA1 could be related to their ability to fluidize the membrane by plotting the EC_50_ value for each compound as a function of the change they induced on exGP^340^ (Figure 5c) and DPH anisotropy (Figure 5d). We found no evidence for such correlation, with coefficients of determination (R^2^) of 0.01 and 0.009 for EC_50_ vs. exGP^340^ and EC_50_ vs. DPH anisotropy, respectively.

**Figure 5.**
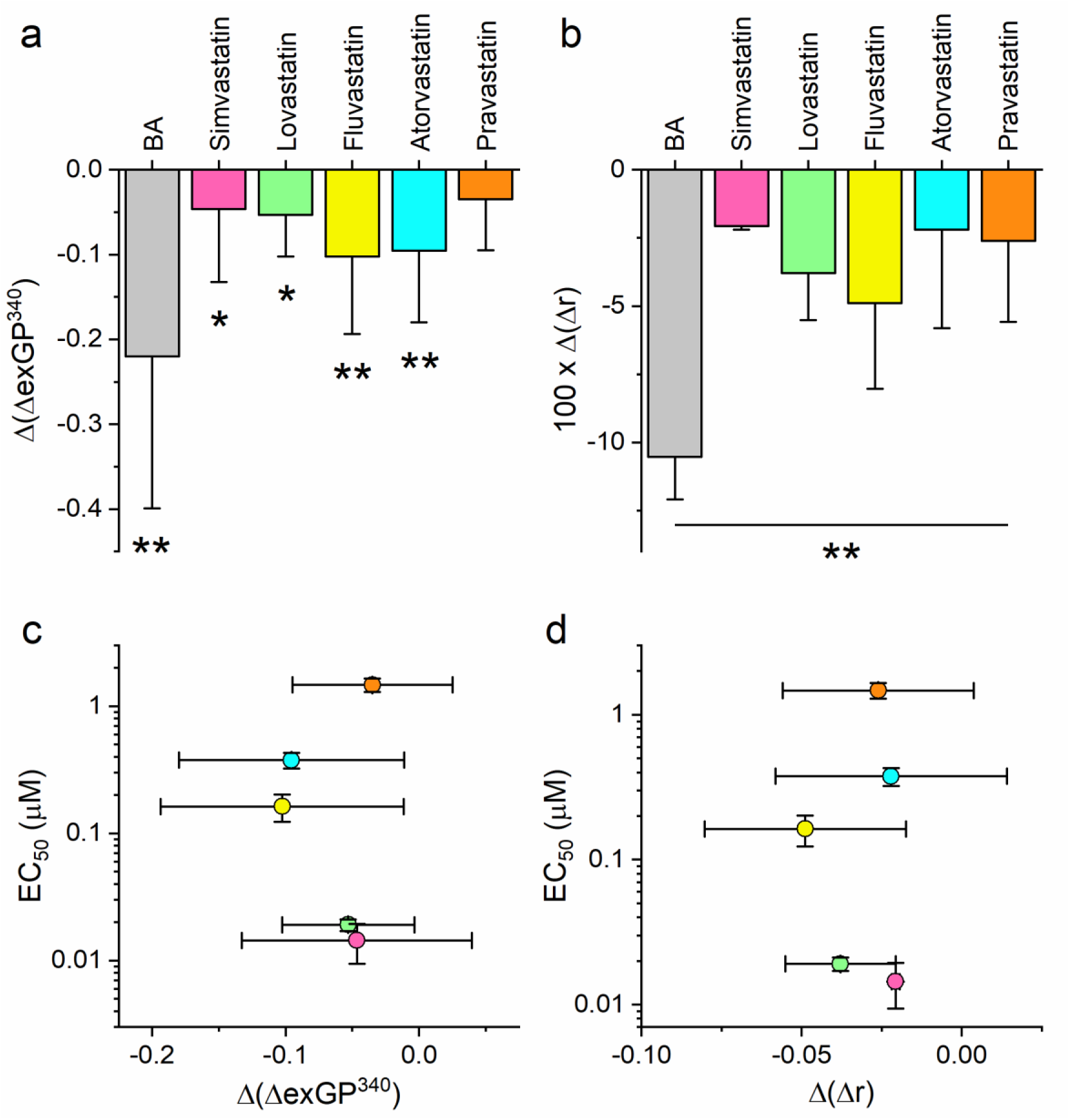
Statins induce cellular membrane fluidization. (a) Changes in exGP^340^ induced by 5 min treatment with 100 µM of statins and 200 mM BA (b) Anisotropy changes induced by 5 min application of 500 µM of statins and 200 mM of BA. All experiments were conducted in CHO-mTRPA1 cells stained with 1.5 µM Laurdan (a) or 10 µM DPH (b) and measured at using FlexStation III. The data are shown as mean ± s.e.m. from at least three experiments. Each point in the scatter plots represents the mean of approximately 20000 cells present in the well of a 96-well plate. Relationship between statins EC_50_ value for TRPA1 activation and change in exGP^340^ (c) or anisotropy (d).

### Activation of mTRPA1 by statins is not mediated by covalent modification of cysteines

Although the statins tested are not electrophiles, it could be hypothesized that they induce the production of an electrophilic compound, which in turn may activate the channel via covalent modification of cysteine residues [28, 29]. We assessed this possibility by testing the effects of statins on a mutant channel in which the residues that largely determine the activation by electrophiles (C622, C642, and C666) are substituted to serine (mTRPA1-3C/S) [22, 28–30]. Application of statins to cells transfected with the mTRPA1-3C/S channel induced Ca^2+^ responses, whose amplitudes were comparable to those of cells transfected with the wild type channel (Figure 6a,b,d). As expected, mTRPA1-3C/S-transfected cells displayed reduced responses to the electrophile AITC (100 μM) (Figure 6a,b,d), whereas the responses to the non-electrophilic agonist, thymol (300 µM) remained unaffected (Figure 6c,d). These results indicate that statin-induced activation of mTRPA1 does not involve the action of electrophilic intermediates.

**Figure 6.**
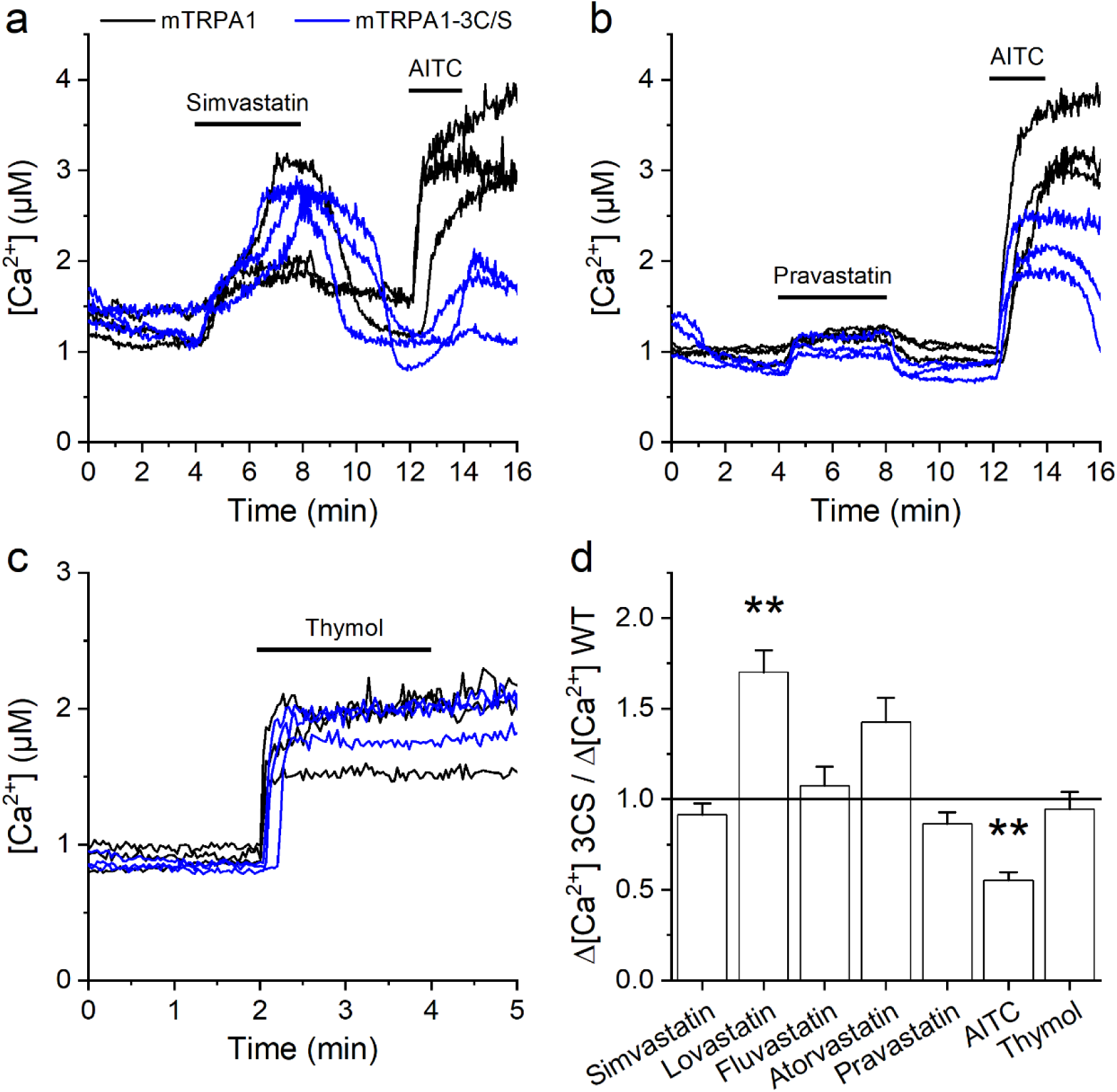
Statin responses are unaltered by a mutation in mTRPA1 N-terminal essential residues. (a-c) Representative traces of [Ca^2+^] responses recorded in cells transfected with the wild-type mTRPA1 or the triple cysteine mutant (mTRPA1-3C/S) in the presence of (a) simvastatin, 10 µM (b) pravastatin, 800 µM, and thymol, 300 µM (c). (d) Average [Ca^2+^] amplitude change evoked by statins in wild-type and mutant mTRPA1. Statins were applied at concentrations 10, 20, 120, 360, and 800 µM for simvastatin, lovastatin, fluvastatin, atorvastatin, and pravastatin, respectively, as well as 100 µM AITC and 300 µM thymol. The error bars represent the standard error of the mean. All experiments were conducted in HEK293T cells transfected with mTRPA1 (n ≥ 112) or mTRPA1-3C/S (n ≥ 111).

### Structural determinants of mTRPA1 activation by statins

As the third hypothesis, we considered whether statins activate TRPA1 by direct interactions with specific sites of the channel protein. To assess this, we sought to identify structural determinants of statin effects employing a strategy based on species-specific effects of non-electrophilic TRPA1 agonists. For this, we took notice that the activation of human and mouse TRPA1 by menthol depends on structural elements in the TM5 and TM6 transmembrane segments that are divergent in the *Drosophila* TRPA1 isoforms [17]. We found that HEK293 cells transfected with dTRPA1(B) responded to simvastatin (100 µM), lovastatin (100 µM), fluvastatin (400 µM) and atorvastatin (1000 µM), but the response amplitudes were significantly smaller than those of mTRPA1 (Figure 7a,b,e). Pravastatin (1500 µM) did not induce detectable responses (Figure 7c,e). Cells transfected with dTRPA1(B) were robustly stimulated by AITC (100 µM) and responded weakly to menthol (1 mM) (Figure 7a-e), as previously reported [17, 31]. As a way to compare the effects of the statins to that of the strong TRPA1 agonist AITC we re-plotted the data in Figure 7e but using the amplitudes of the responses to the statins normalized to the amplitude of the subsequent response to AITC in each cell (Figure 7f). These normalized data show that the statins induced much weaker relative responses in dTRPA1(B) than in mTRPA1, indicating for a reduction of the sensitivity to the statins in the *Drosophila* isoform.

**Figure 7.**
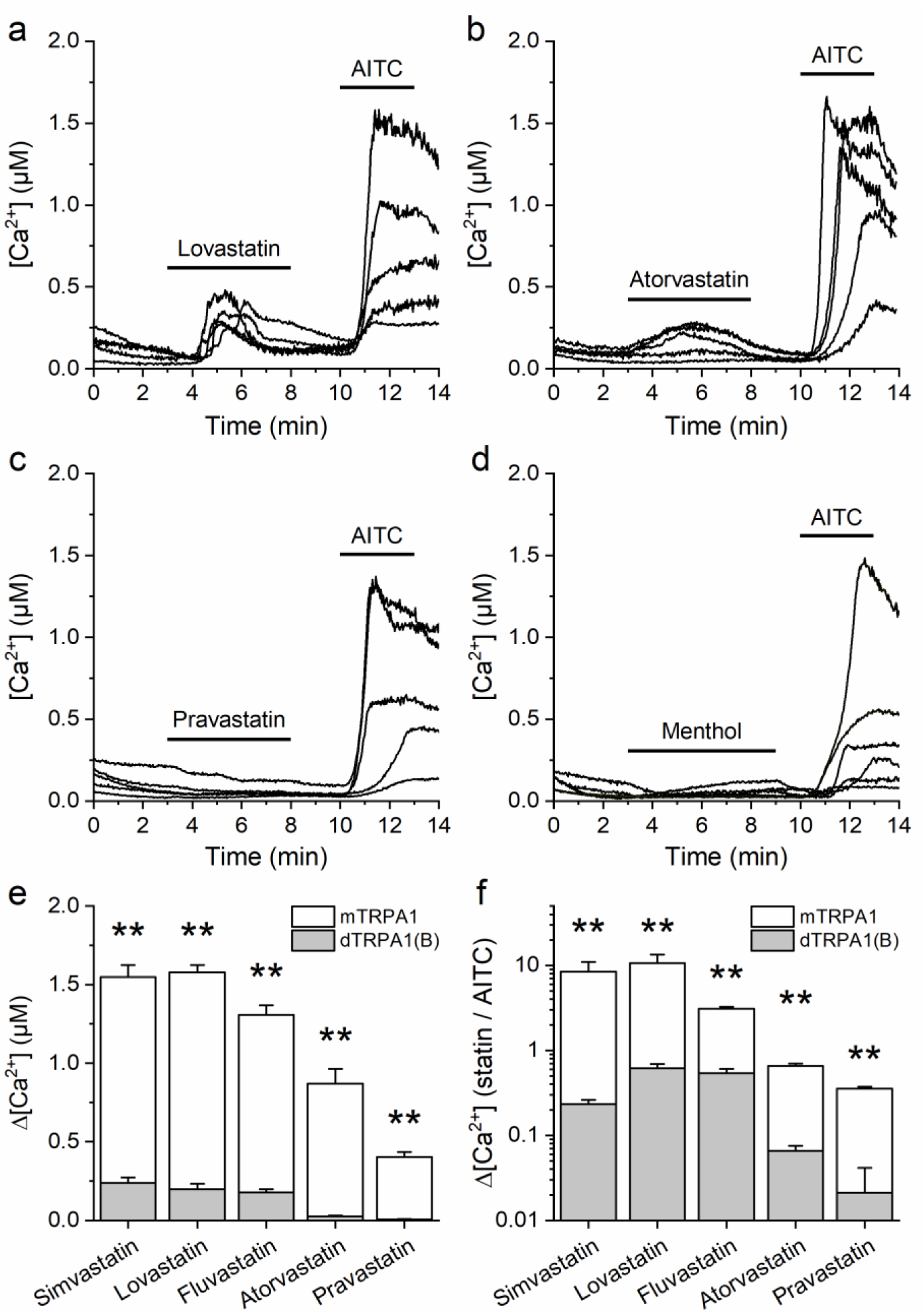
Statin responses are reduced in cells transfected with the dTRPA1(B) variant. (a-d) Representative traces of [Ca^2+^] responses recorded in cells transfected with the dTRPA1 variant B in the presence of (a) lovastatin, 100 µM, (b) atorvastatin, 1000 µM, (c) pravastatin, 1500 µM, and menthol, 100 µM (d). (e) Comparison of the average [Ca^2+^] amplitude change evoked by statins in cells transfected with mTRPA1 (white bar) and dTRPA1(B) (gray bar). (f) Average [Ca^2+^] amplitude change evoked by statins normalized to the amplitude of the subsequent response to AITC in dTRPA1(B) (gray bar) and mTRPA1 (white bar). Statins were applied at concentrations of 100, 100, 400, 1000, and 1500 µM for simvastatin, lovastatin, fluvastatin, atorvastatin, and pravastatin, respectively. The error bars represent the standard error of the mean. All experiments were conducted in HEK293T cells transfected dTRPA1(B) (n ≥ 67) or with mTRPA1 (n ≥ 94).

These results suggest that the mechanisms underlying the activation of TRPA1 by statins and menthol share similar structural determinants. To assess this possibility, we performed molecular docking simulations (9 per compound) using a model of mTRPA1 generated from the cryo-EM human TRPA1 structure [32]. We found that menthol and all statins mainly docked to two non-overlapping putative binding sites formed by amino acid residues located in the TM4, TM5 and TM6 segments (Figure 8a-f). Specifically, site 1 is near a kink in the middle of TM5, a region that is in close proximity to TM6. Site 2 is formed by amino acids of the end of TM4 and the proximal part of TM5. The estimated binding energies of the statins were found to be larger than those of menthol (Figure 8g). Binding site 1 exhibited a higher chance of occupancy by all compounds, particularly for the more hydrophilic statins, atorvastatin and pravastatin, which were primarily located at this site (Figure 8h).

**Figure 8.**
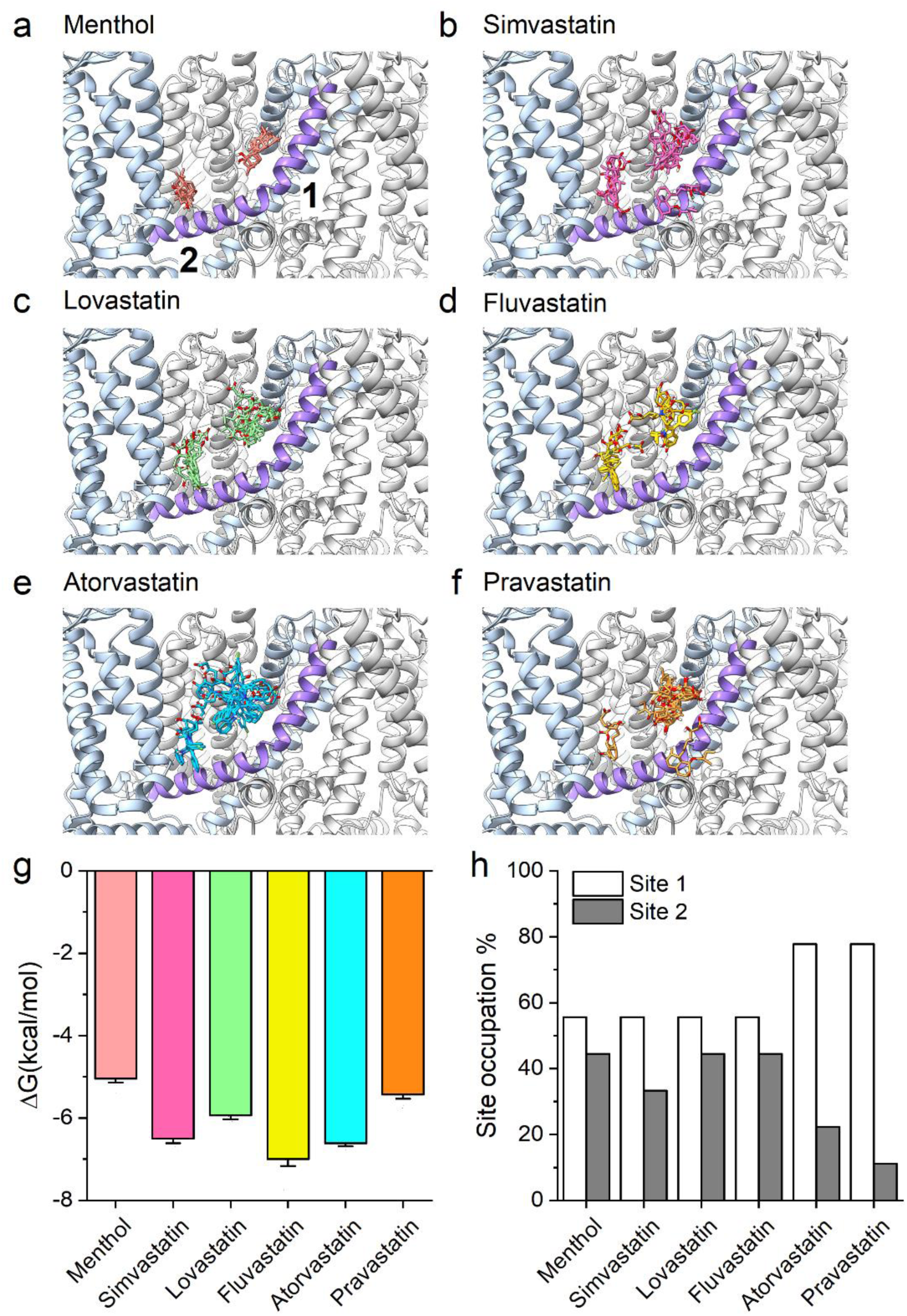
Model of statins binding to mTRPA1. (a-f) Model of the mTRPA1 channel presenting localization of TM5 (colored purple) and (a) menthol, (b) simvastatin, (c) lovastatin, (d) fluvastatin, (e) atorvastatin, and (f) pravastatin docking. 1 and 2 represent possible binding sites. (g) Calculated energies of menthol and statins binding to the docking sites in the mTRPA1 channels (n = 9). (h) Percentage of binding site 1 (white bar) and binding site 2 (gray bar) occupation by menthol and statins.

To validate the results of the docking simulations, we tested the effects of menthol (100 µM), simvastatin (10 µM), lovastatin (20 µM), fluvastatin (120 µM) and atorvastatin (360 µM) on mTRPA1 mutants in which single amino acids were mutated to the corresponding ones in dTRPA1(B). We did not evaluate the responses to pravastatin because they were too small to be reliable for comparisons. We first tested the effect of an alanine substitution of residue F950, which is located in TM6 and found to interact at site 1 with simvastatin, fluvastatin and atorvastatin in the docking simulations. HEK293 cells transfected with the F950A mutant showed reduced responses to precisely those statins, but preserved responses to AITC, lovastatin and enhanced response to menthol (Figure 9b,e). We tested as well the implication of residues S876 and T877 in TM5, also located in site 1, and that were associated with the activation by menthol [17]. From site 2, we tested the TM4 residue F843, whose corresponding one in the human orthologue, Y840, was suggested to be part of the binding site of the non-electrophilic activators delta-9-tetrahydrocannabinol (THC) and cannabidiol (CBD) [20]. We found that mutants S876V, T877L and F843A displayed significantly smaller responses to menthol and all statins (Figure 9 b-e). However, S876V also responded very weakly to AITC, suggesting for a more general role of this residue in channel activation and/or membrane expression. The data also shows that, except for atorvastatin, the mutant F843A in site 2 displayed smaller responses than the mutants F950A and T877L in site 1 (Figure 9e).

**Figure 9.**
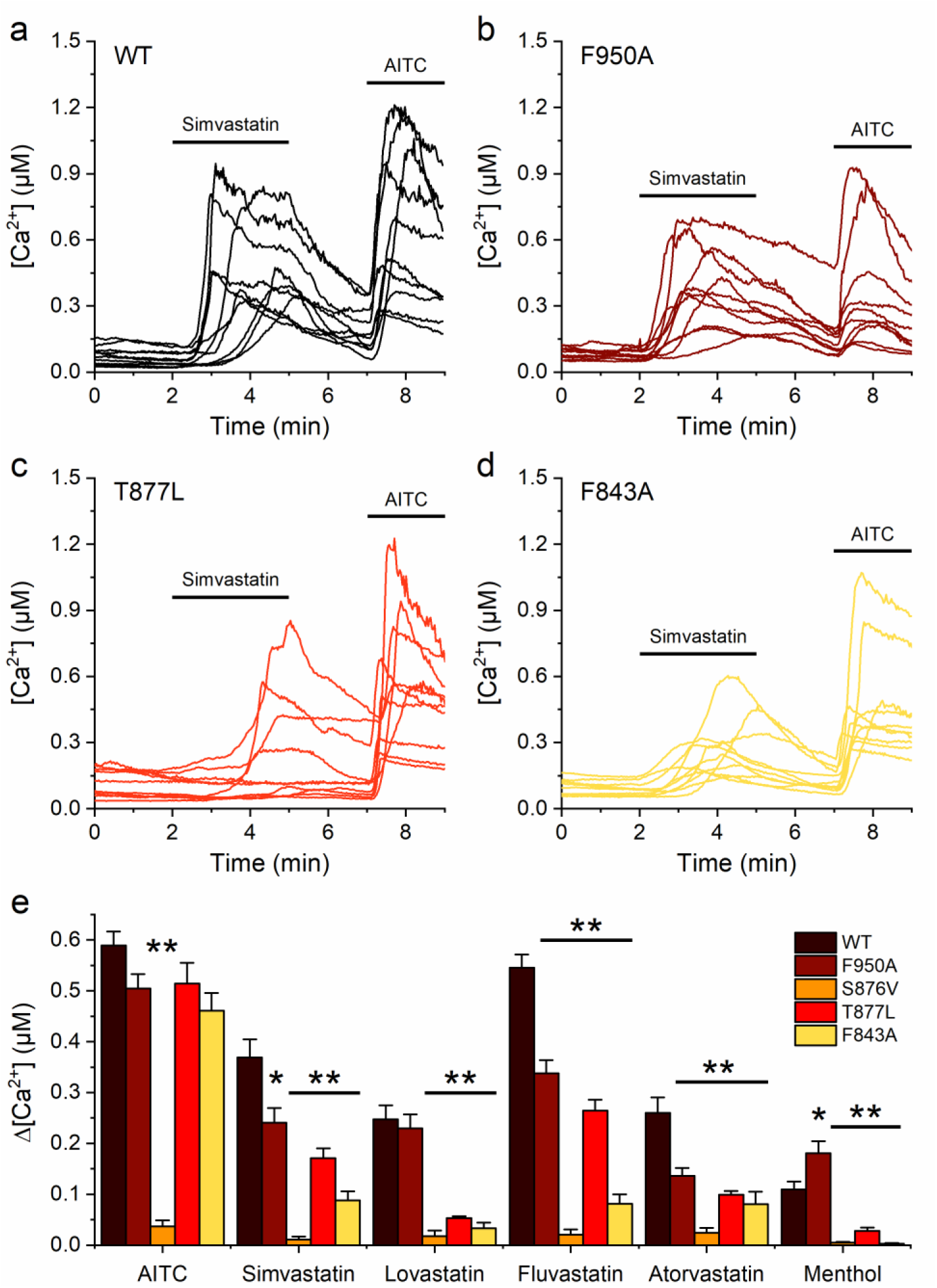
Single-point mutations in mTRPA1 binding sites affect responses to statins. (a-d) Representative traces of [Ca^2+^] responses recorded in HEK293T cells transfected with mTRPA1 (a) and single-point mTRPA1 mutants: F950A (b), T877L (c), and F843A (d), in the presence of simvastatin (10 µM) and AITC (100 µM). (e) Average [Ca^2+^] amplitude change evoked by statins in wild-type and mutant mTRPA1. Statins were applied at concentrations of 10, 20, 120, 360, and 800 µM for simvastatin, lovastatin, fluvastatin, atorvastatin, and pravastatin, respectively, as well as 100 µM menthol. Error bars represent the standard error of the mean. All experiments were conducted in transfected HEK293T cells, with n ≥ 50 for all conditions presented.

The residue T877 belongs to a stretch that was speculated to contribute to the binding of the inhibitor A-967079 to hTRPA1 [33]. Thus, it was interesting to test whether this could be confirmed experimentally in mTRPA1 and to determine the effects of A-967079 on the agonist actions of menthol and the statins, which are partly mediated by this residue. We found that A-967079 strongly decreased responses to AITC in cells transfected with WT mTRPA1 (p < 0.0001), but had a marginal inhibitory effect in cells transfected with the T877L mutant (p = 0.0376; Figure 10a,c). A-967079 inhibited the WT responses to the statins and menthol, but surprisingly, it potentiated the responses of the T877L mutant to these compounds (Figure 10b,c). These data are better visualized by plotting the ratio of the average amplitudes of the responses to the agonists determined in the presence and in the absence of A-967079 (Figure 10d). Two observations can be done in this plot: one, that A-967079 inhibits more the response to AITC than the responses to the statins and menthol, and two, that the effects of A-967079 seem to vary across the different agonists in the same proportion for the WT and T877L mutant. To verify this, we plotted the effects on the mutant versus those on the WT, and indeed found that these are highly correlated across agonists (R^2^ = 0.97). Taken together, these data indicate that the residue T877 is implicated in the inhibitory action of A-967079 on WT mTRPA1, and demonstrate that A-967079 is able to interact with the T877L mutant. Moreover, the effect of A-967079 depends on the nature of the agonist, in a way that the less it inhibits the effect of the agonist on the WT channel, the more it enhances the effect of the agonist in the T877L mutant.

**Figure 10.**
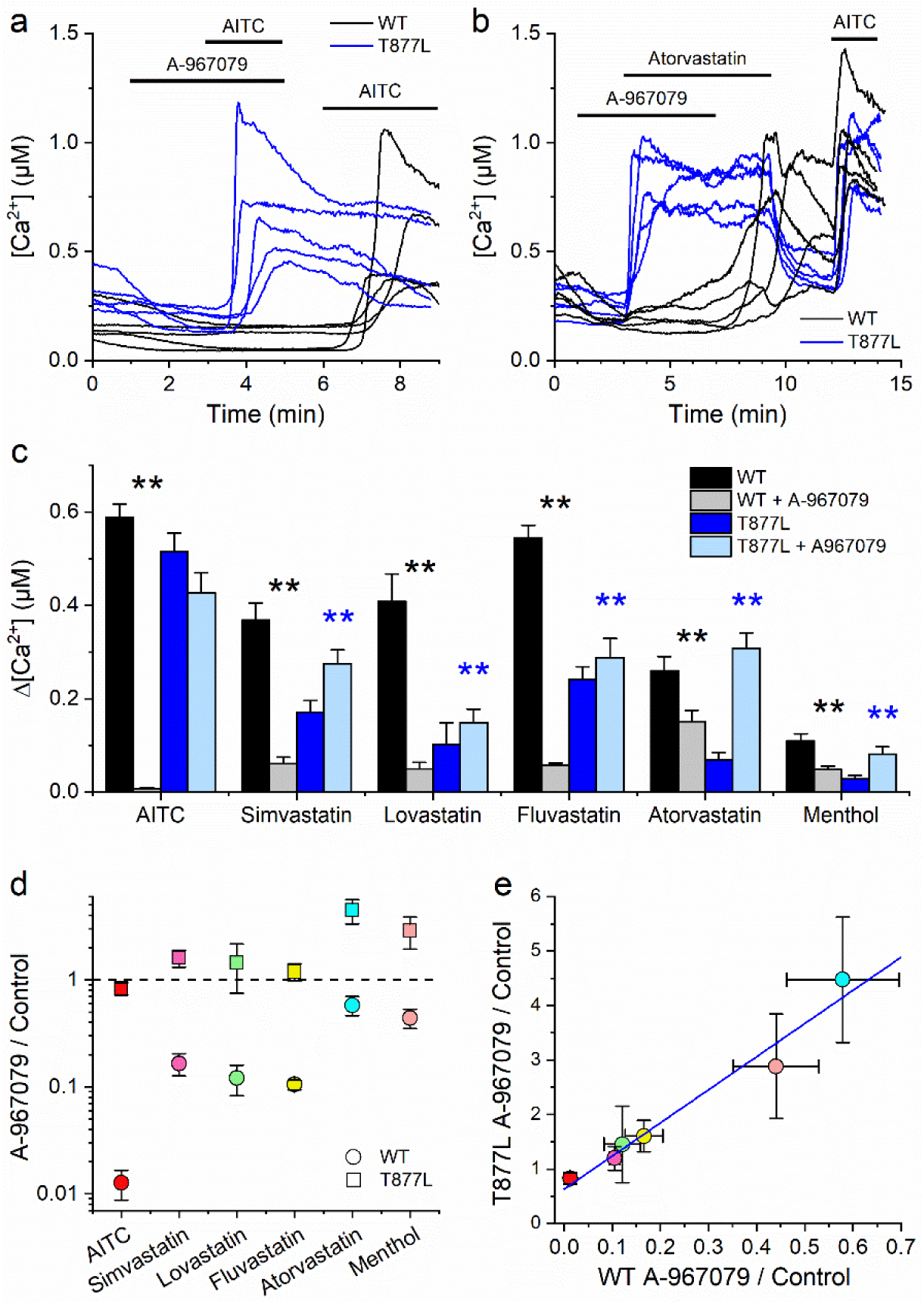
Binding A-967079 in the binding pocket of the T877L mutant induces potentiation of statin response. (a and b) Representative traces of [Ca^2+^] responses recorded in HEK293T cells transfected with the mTRPA1 (black trace) and single point mTRPA1 mutant T877L (blue trace) in the presence of (a) AITC (20 µM) and (b) atorvastatin (400 µM) with and without a channel antagonist A-967079 (10 µM). (c) Average [Ca^2+^] amplitude change evoked by AITC, statins, and menthol in mTRPA1 (black and gray bar; without or with inhibitor) and T877L mutant (blue and light blue bar; without or with inhibitor). (d) The ratio of the average amplitudes of the responses to AITC, statins, and menthol determined in the presence and in the absence of A-967079 in mTRPA1 (circles) and mutant T877L channels (squares). (e) Correlating the effects on the T877L mutant versus those on the WT channel with or without A-967079. Statins were applied at concentrations of 10, 20, 120, and 360 µM for simvastatin, lovastatin, fluvastatin, and atorvastatin, respectively, as well as 100 µM menthol (n ≥ 100 for all presented conditions).

## Discussion

Previous work has established that TRPA1 activation and membrane expression are sensitive to manipulations of plasma membrane cholesterol [4, 5, 34], suggesting that TRPA1 function can be altered by cholesterol-reducing drugs. Indeed, recent studies showed that cyclodextrin derivatives reduce TRPA1 activation [35, 36]. However, the broad chemical sensitivity of TRPA1 [2, 3] and the conflicting results of previous studies on the effects of statins on this channel [15, 16], prompted us to (re-)evaluated the actions of five of such compounds on hTRPA1 and mTRPA1.

We found that five statins (simvastatin, lovastatin, fluvastatin, atorvastatin, and pravastatin) activate recombinant mouse and human TRPA1 channels in a concentration-dependent and reversible manner. Experiments performed in a native expression system, mouse DRG neurons, showed that TRPA1-expressing neurons responded to statins and that HC-030031 largely inhibited these responses. Our results on simvastatin agree with a recent report showing activation of hTRPA1 at micromolar concentrations, as well as TRPA1-independent responses in mouse DRG neurons [16]. The effects of the other statins, however, were largely dependent on TRPA1 at the concentrations that we tested, close to the EC_50_ values for mTRPA1. These data show that TRPA1 is a direct effector of statins, rather than just an element recruited downstream of TRPV1 activation as previously suggested [15]. These discrepancies may be explained by the distinct statin application methods, i.e, acute ([16] and present results) vs. 2 h pretreatment [15]. Indeed, because statins can inhibit cholesterol synthesis, leading to a significant reduction of available lipid rafts in the membrane where TRPA1 is located, long-term exposures may reduce expression and sensitivity of TRPA1, as previously observed upon pretreatment with MCD or SMase [4, 5].

The relevance of our results may be sought in several directions. The first rises from noting that lovastatin is a natural compound, belonging to a family of secondary metabolites produced by fungi such as *Aspergillus terreus* [37]. Future studies may address whether TRPA1 activation by statins has any ecological significance, as previously proposed for many plant-derived agonists of this channel [38]. The second potential relevance relates to the therapeutic context, and specifically to the question of whether statin concentrations used for the treatment of hypercholesterolemia reach the supra-micromolar levels required for TRPA1 activation. This does not seem to be the case for common dosage schemes, as the therapeutic levels in human plasma are reported in the range from 1 to 15 nM, with peak concentrations at up to 50 nM [39]. On the other hand, the activation of sensory TRP channels by statins might be relevant for the interpretation of results of in vitro and in vivo rodent experiments in which high micromolar doses of these drugs are administered (1 to 50 µM). Moreover, some clinical studies employing maximum tolerated doses of lovastatin (25 mg/kg, daily [40]) or simvastatin (7.5 mg/kg, twice daily [41]) reported maximal plasma concentrations between 0.1-3.9 μM and 0.08-2.2 μM, respectively, which are at the low end of the effects on TRPA1, TRPV1 and TRPM8 ([16] and present results). Nevertheless, a previous suggestion that activation of these channels is clinically relevant for the reported analgesic and anti-inflammatory actions of statins [16] should be taken with caution. Instead, this could be related to a decrease of membrane cholesterol itself, which is known to decrease TRPA1 membrane expression and activation [4, 5, 34].

Finally, having identified another family of non-electrophilic compounds as TRPA1 agonists allowed us to obtain valuable insight into the puzzling ability of this channel to be activated by a myriad of chemically-unrelated agonists [2, 3]. We tested three potential mechanisms, the first one consisting on the induction of mechanical perturbations in the plasma membrane, as proposed for trinitrophenol [42], primary alcohols [43], bacterial lipopolysaccharides (LPS) [24] and alkylphenols [22]. Our results with two independent reporting dyes show that all statins induced disordering of the plasma membrane. However, these effects were not predictive of the EC_50_ to activate mTRPA1, contrasting to what we have previously shown for LPS [24] and alkylphenols [22]. Similarly, we found no evidence of mTRPA1 activation by statins through covalent modification of the channel, as the responses to these compounds where not decreased in a triple cysteine mutant channel that has reduced sensitivity to electrophilic compounds [28–30].

Finally, we considered the possibility that statins may activate TRPA1 by direct interactions. The finding that the dTRPA1(B) isoform responds weakly to all statins pointed to the implication of structural elements in the TM5 and TM6 transmembrane segments, as previously reported for the non-electrophilic agonist menthol [17]. This suspicion was further supported by docking simulations indicating that menthol and all statins interact with two distinct bindings sites centered around TM5 in the middle-lower pore domain but also involving residues of TM4 and TM6.

Site 1 corresponds to a previously proposed menthol-binding pocket, with critical implication of TM5 residues S876 and T877 [17]. This site overlaps with a large tertiary binding pocket of antagonists A-967079 [32, 44] and GDC-0334 [45]. It has been proposed that at least 20 different amino acids of the TM5 helix, the TM6 helix, the pore helix 1, and the TM4-TM5 linker are involved in that binding pocket. Site 2 overlaps with the binding pocket of the non-covalent agonists GNE551 [19], β-eudesmol [18], THC and CBD [20]. Menthol, simvastatin, lovastatin and fluvastatin bound to site 1 in 55% of the simulations, but atorvastatin and pravastatin showed more preference for this site (80%). The results with single-point mutants F950A and T877L (site 1) and of F843A (site 2) on the responses to menthol are difficult to interpret because the two mutations in site 1 produced opposite effects, i.e., enhancement of the response for F950A and decrease for T877L. This may be related to the bimodal effect of menthol on mTRPA1, from which it is expected that mutation of some residues disrupts more the agonist than the antagonist effect, whereas this can be the opposite for mutation of other residues. In contrast, the mutations in both putative binding sites resulted in consistent reductions of the agonist actions of all statins. This probably reflects the fact that, as a difference from menthol, none of the 5 statins tested displayed a bimodal effect (i.e., decreasing effects at high concentrations or rebound responses upon washout) [46]. Yet, it was also apparent that the impact of the mutation of given residues in sites 1 and 2 was different across the distinct statins. For instance, mutation F950A impacted the responses to simvastatin, fluvastatin and atorvastatin, but not that of lovastatin, which is coherent with the results of docking simulations showing that the latter does not engage residue F950. Taken together, these results provide evidence for 2 binding sites, whose relative importance for the agonist effect is, as expected, molecule-specific. Future research may help testing whether, as our docking simulations suggest, these differences could be related to the distinct relative occupation, binding energies and amino acids that each statin engages in the two binding sites.

Further evidence of the presence of these regulatory sites in mTRPA1 was found with the known inhibitor A-967079. Firstly, this compound largely abrogated the response to the electrophilic agonist AITC, but had weaker and varied effects across different statins and menthol. Secondly, mutation T877L in site 1 strongly reduced the inhibitory action of A-967079 on the responses to AITC, but in sharp contrast, did not affect the activation by lovastatin and fluvastatin and actually enhanced the effects of simvastatin, atorvastatin and menthol. These results show that the inhibitor competes with the statins and menthol for site 1 but not with AITC, which again is consistent with the latter covalently modifying amino acid residues located somewhere else in the channel.

Finally, another element influencing the agonist activity of statins is their ability to partition in the plasma membrane and from there reaching the binding sites in TRPA1. From that perspective, the statins we tested manifest distinct distribution in lipid phases and membranes. According to studies in artificial lipid micelles and membranes, simvastatin and lovastatin locate well within the hydrophobic, hydrocarbon tails of the lipids, fluvastatin also penetrates into the membrane but with a shallower incorporation, atorvastatin penetrates membranes but its polar moiety remains at the interfacial region of the bilayer, and pravastatin interacts only at the surface and does not permeate the membrane [47]. These findings are in line with the lipid/water distribution coefficient of the statins, which show the order: simvastatin (4.4), lovastatin (3.91), fluvastatin (1.75), atorvastatin (1.53) and pravastatin (−0.47). In fact, the EC_50_ and the distribution coefficient show a very strong inverse correlation, with a Pearson’s R value of −0.99 and an R^2^ = 0.98 (not shown). Interestingly, a similar result was previously considered to support the hypothesis that alkylphenols activate TRPA1 by inducing mechanical perturbations in the plasma membrane, as higher membrane partition may enhance the ability of compounds to modify the local lipid environment around the channel. However, as discussed above, we found no evidence for such an effect with the statins. Thus, our present findings may suggest that also alkylphenols interact with the putative TRPA1 binding sites we describe here, a possibility that should be checked in future experiments.

In summary, our results demonstrate that different statins activate human and mouse TRPA1 channel in a concentration-dependent and reversible manner. The underlying mechanism seems to involve binding to two tertiary pockets that can accommodate various compounds, including agonists and inhibitors simultaneously. The identification of two distinct agonist binding sites may help explaining how TRPA1 is able to respond to a large variety of non-electrophilic compounds, while the finding of competitive interactions at one of these sites may help guide the development of agonist-specific antagonists of therapeutic relevance.

## Methods

### Cell culture

Human embryonic kidney (HEK293T) cells from the European Collection of Cell Culture (Salisbury, UK) were grown in Dulbecco’s modified Eagle’s medium (DMEM) containing 10% (v/v) fetal calf serum, 2 mM L-glutamine, 2 units/ml penicillin and 2 mg/ml streptomycin (Gibco/Invitrogen, Carlsbad, CA, USA) at 37 °C in a humidity-controlled incubator with 10% CO_2_. HEK293T cells were transiently transfected with the pCAGGS-IRES-GFP vector [48] encoding human, drosophila (variant B) TRPA1, or pCAGGSM2-mCherry encoding mTRPA1 or single point mutants F843A, T877L, S876V, F950A. Transfected cells were reseeded after approximately 16 h on poly-L-lysine-coated (0.1 mg/ml) 18 mm glass coverslips with a thickness of 0.13 - 0.16 mm (Gerhard Menzel GmbH) for ratiometric intracellular Ca^2+^ imaging.

Chinese hamster ovary (CHO-K1) cells from the American Type Culture Collection were grown in DMEM containing 10% fetal bovine serum, 2% glutamax (Gibco/Invitrogen), 1% non-essential amino acids (Invitrogen), and 200 µg/ml penicillin/streptomycin at 37 °C in a humidity-controlled incubator with 5% CO_2_. As TRPA1 expression system, we used CHO-K1 cells stably transfected with mouse TRPA1 (CHO-TRPA1) in an inducible system [49]. For patch-clamp measurements TRPA1 expression was induced by incubation with 1 μg/ml tetracycline for at least 12 h.

### Isolation and culture of DRG neurons

Mouse dorsal root ganglion (DRG) neurons were cultured using an adapted protocol of a previously published method [50]. For each series of experiments, 1 to 2 adult mice (10-12 weeks, male and female) were killed by gradual exposure to CO_2_ and the DRGs along the spine were bilaterally excised under a dissection microscope. The ganglia were kept in Dulbecco’s phosphate buffered saline (Gibco, Grand Island, NY, USA) during the isolation procedure and placed in a 2 ml tube containing 800 ul 10% fetal calf serum Neurobasal A medium (basal medium) and later incubated at 37 °C in a mix of 2 mg/ml collagenase (Gibco) and 2.5 mg/ml dispase (Gibco) for 60 min. Digested ganglia were gently washed twice with basal medium and once with B27-supplemented Neurobasal A medium (Invitrogen) containing 2 mM Glutamax, 2 ng/ml glial cell line-derived neurotrophic factor (Invitrogen), 10 ng/ml neurotrophic factor 4 (PeproTech, Rocky Hill, NJ, USA) and 1% PenStrep (complete medium). Neurons were mechanically dissociated by mixing with syringes fitted with increasing needle gauges, centrifuged through bovine serum albumin (BSA) gradient (from 16% to 0%) in order to filter out the debris, and distributed in 6 poly(l-lysine)/laminin-coated glass chambers (Fluorodish; World Precision Instruments, Hitchin, Hertfordshire, UK) and cultured for 12–18 h at 37 °C in complete medium.

### Ratiometric intracellular Ca^2+^ imaging

For intracellular Ca^2+^ imaging experiments, cells were incubated with 2 µM Fura-2 AM (Biotium, Hayward, CA, USA) for 30 min at 37 °C in a humidity-controlled incubator. Fluorescence was measured with alternating excitation at 340 and 380 nm using a monochromator-based imaging system consisting of an MT-10 illumination system (Tokyo, Japan) and Cell^M^ software from Olympus. All experiments were performed at 25 °C using a standard Krebs solution containing (in mM) 150 NaCl, 6 KCl, 1.5 CaCl_2_ x 2H_2_O, 1 MgCl_2_ x 6H_2_O, 10 HEPES, 10 glucose (pH 7.4 with NaOH). Fluorescence intensities were corrected for background signal and presented as the ratio F340/F380 from which intracellular Ca^2+^ was calculated as described previously [51]. Ca^2+^ ratios were calculated as the difference between the maximum and basal values of responding cells during the stimulus period. Only cells that responded to the positive control, AITC, at the experiment’s end were considered TRPA1-expressing cells. All experiments were performed on at least three independent coverslips. Data points were analyzed and presented as means ± s.e.m of the given number (n) of individual experiments OriginPro 2018 (OriginLab Corporation). A Hill function of the form fitted the experimental concentration dependences:

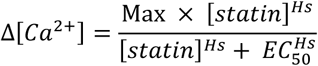

where Max stands for the maximal increase in [Ca^2+^]; [statin] is the statin concentration; EC_50_ is the effective concentration and *H*_S_ is the corresponding Hill coefficient.

### Electrophysiology

Whole-cell patch-clamp recordings were performed with an EPC-10 amplifier and the PatchMasterPro Software (HEKA Elektronik, Lambrecht, Germany). Currents were sampled at 20 kHz and digitally filtered at 2.9 kHz. The internal solution contained (in mM): 156 CsCl, 1 MgCl_2_, 10 HEPES and 10 BAPTA (pH 7.2 with CsOH). The extracellular solution contained (in mM): 150 NaCl, 2 CaCl_2_, 1 MgCl_2_ and 10 HEPES (pH 7.4 with NaOH). The patch pipette resistance was between 2 and 4 MΩ when filled with pipette solution. Solutions were applied via a gravity-based multi-barrel pipette tip with a single outlet, allowing for a solution exchange around the cell in 2-4 s. Valves were operated manually via an electronic gear-box by the experimenter. Currents were elicited at intervals of 2 s with a 500 ms ramp protocol ranging from −100 mV to +100 mV. Data were analyzed using IgorPro 6.2 (WaveMetrics, USA) and OriginPro 2018 (OriginLab Corporation, USA). All data sets were tested for normality, and a Grubbs test was used to identify outliers excluded from the analyses. A one-sample *t*-test was used to determine the significance of current increases.

### Fluorescent measurements using Laurdan and DPH

A 0.8 mM stock solution of Laurdan (6-dodecanoyl-N,N-dimethyl-2-naphthylamine) was prepared in methanol (Sigma-Aldrich, Bornem, Belgium). Next, 1.5 µM Laurdan was prepared in Krebs solution (see above) to stain CHO-mTRPA1 cells (1×10^6^ cells/ml) for 30 min at 37 °C. After incubation, cells were washed with Krebs, and suspensions of 100 μl were aliquoted into flat-bottom 96-well microtiter plates (Greiner Bio-One). Steady-state Laurdan fluorescence measurements were performed using a FlexStation 3 Benchtop Multi-Mode Microplate Reader and the SoftMax Pro Microplate Data Acquisition & Analysis Software (Molecular Devices). Generalized polarisation (GP) values were calculated using emission values at 440 nm and 490 nm, and excitation at 340 nm, according to the formula exGP^340^ = (I_440_ - I_490_)/(I_440_ + I_490_) [25].

A 1 mM stock solution of DPH (Sigma-Aldrich) was prepared in dimethyl sulfoxide (DMSO; Sigma-Aldrich), and a working solution of 10 µM was prepared in Krebs. Cells were stained as described above for Laurdan. DPH anisotropy (r) was monitored with an excitation wavelength of 365 nm and emission wavelength of 430 nm using a Flexstation 3 Benchtop Multi-Mode Microplate Reader and the SoftMax Pro Microplate Data Acquisition & Analysis Software (Molecular Devices). A linearly polarised excitation beam was generated by a vertical polariser that excites DPH with transition moments aligned parallel to the incident polarisation vector. The resulting fluorescence intensities in both parallel (I_VV_) and perpendicular (I_VH_) directions to that of the excitation beam were recorded, and the fluorescence anisotropy can be calculated by: r = (I_VV_ - G·I_VH_)/(I_VV_ + 2G·I_VH_), where G = I_HV_/I_HH_. The relation between anisotropy (r) and polarisation (P) was determined as P = 3r/(2 + r), and the membrane fluidity was calculated as the inverse of fluorescence polarisation (1/P) [52].

### Simulations of docking of menthol and statins on mTRPA1

The structure of mouse TRPA1 was built by homology modeling based on the experimentally determined structure of its human homolog (PDB code: 3J9P) [32], using the program Modeller [53]. The software Autodock-Vina [54] was used to predict the possible binding modes of menthol and statins to the TM5 transmembrane segments.

### Data and statistical analysis

The key conclusions of the manuscript are supported by experimental recordings of n ≥ 5 individual cells per group for patch-clamp experiments, n ≥ 50 for ratiometric Ca^2+^ imaging, and n ≥ 3 wells for membrane perturbation evaluation, with the exact group numbers specified in the Figure legends and main text. We used the following parameters for the statistical analysis: a 20% difference between groups, a standard deviation of 5%, a significance level of 0.05, and a power of 0.95. Group sizes for patch-clamp experiments vary due to the complexity of the technique, leading to differences in protocol success rates.

Data recordings were randomized by random selection of CHO and HEK293T cells expressing TRPA1 and DRG neurons on each individual cover slip, but operators were not blinded during data collection and analysis due to the experimental procedure and the range of used concentrations needed for each of the compounds. The electrophysiological recordings are operator-independent and free of subjective interpretation. Ratiometric Ca^2+^ imaging data was analyzed from all cells in the slide with a prewritten analysis script (Origin; OriginLab Corporation, USA), thereby further canceling any operator bias. Data present in this manuscript was analyzed using Cell^M^ software (Olympus), Excel (Microsoft), IgorPro 6.2 (WaveMetrics, USA), WinASCD (Guy Droogmans, Leuven, Belgium), SoftMax Pro Microplate Data Acquisition & Analysis Software (Molecular Devices), GraphPad Prism version 5.01 (GraphPad Software, Inc.) and Origin (OriginLab Corporation, USA). Origin was further used for statistical analysis and data display. In the Figures, asterisks indicate statistically significant differences (*, p < 0.05; **, *p* < 0.01).

### Materials

Simvastatin (CAS Registry No.: 79902-63-9; Purity ≥ 98%), lovastatin (CAS Registry No.: 75330-75-5; Purity ≥ 98%), fluvastatin (CAS Registry No.: 93957-54-1; Purity ≥ 98%), atorvastatin (CAS Registry No.: 357164-38-6; Purity ≥ 98%), pravastatin (CAS Registry No.: 81093-37-0; Purity ≥ 98%) were purchased from Cayman Chemical (Michigan, USA) and were prepared in DMSO as a stock solution (50 mM) and kept at −20 °C. Allyl isothiocyanate (AITC; CAS Registry No.: 57-06-7; Purity ≥ 95%) from Sigma-Aldrich (Bornem, Belgium) stock in ethanol (100 mM). Cinnamaldehyde (CA; CAS Registry No.: 14371-10-9; Purity ≥ 99%) from Sigma-Aldrich (Bornem, Belgium) stock in ethanol (100 mM). Capsaicin (caps; CAS Registry No.: 404-86-4; Purity ≥ 95%) from Sigma-Aldrich (Bornem, Belgium) stock in ethanol (20 mM). Thymol (CAS Registry No.: 89-83-8; Purity ≥ 98.5%) from Sigma-Aldrich (Bornem, Belgium) stock in ethanol (1 M). HC-030031 (CAS Registry No.: 349085-38-7; Purity ≥ 98%) from Sigma-Aldrich (Bornem, Belgium) stock in DMSO (100 mM). A-9607079 (CAS Registry No.: 1170613-55-4; Purity ≥ 98%) from Tocris (Bio-Techne Ireland Limited, Ireland) stock in ethanol (10 mM). Laurdan (6-dodecanoyl-N,N-dimethyl-2-naphthylamine; CAS Registry No.: 74515-25-6; Purity ≥ 97.0%) from Sigma-Aldrich (Bornem, Belgium) stock in methanol (0.8 mM). DPH (1,6-Diphenyl-1,3,5-hexatriene; CAS Registry No.: 1720-32-7; Purity ≥ 98%) from Sigma-Aldrich (Bornem, Belgium) stock in DMSO (1 mM). All stock solutions were kept at −20 °C, and dilutions to the desired concentrations were freshly prepared from the stock solutions in the experimental solutions.

## Author Contributions

Conception and design: KT, JBS; Acquisition of data: JBS, AM, KH, AT; Analysis and interpretation of data: KT, AM, KH, AT, JBS; Drafting of the manuscript: KT, AM, JBS; Critical revision of the manuscript for important intellectual content: KT, AM, KH, AT, JBS. All listed authors approved the final version of the manuscript and agreed to be accountable for all aspects of the work to ensure that questions related to the accuracy or integrity of any part of the work are appropriately investigated and resolved. All of the above-listed authors qualify for authorship, and all persons qualifying for authorship are listed above.

## Acknowledgments

We thank Melissa Benoit for the technical support and the members of the LICR for helpful discussions. The CHO-mTRPA1 cell line was kindly provided by Dr. Ardem Patapoutian (The Scripps Research Institute, USA). The research was supported by grants from the Research Council of the KU Leuven (C14/18/086) and the Fund for Scientific Research Flanders (FWO: G0D0417N and G0B2219N. J.B.S is a Postdoctoral Fellow of the FWO (12W6122N).

---

**Figure 2 Supplement 1.**
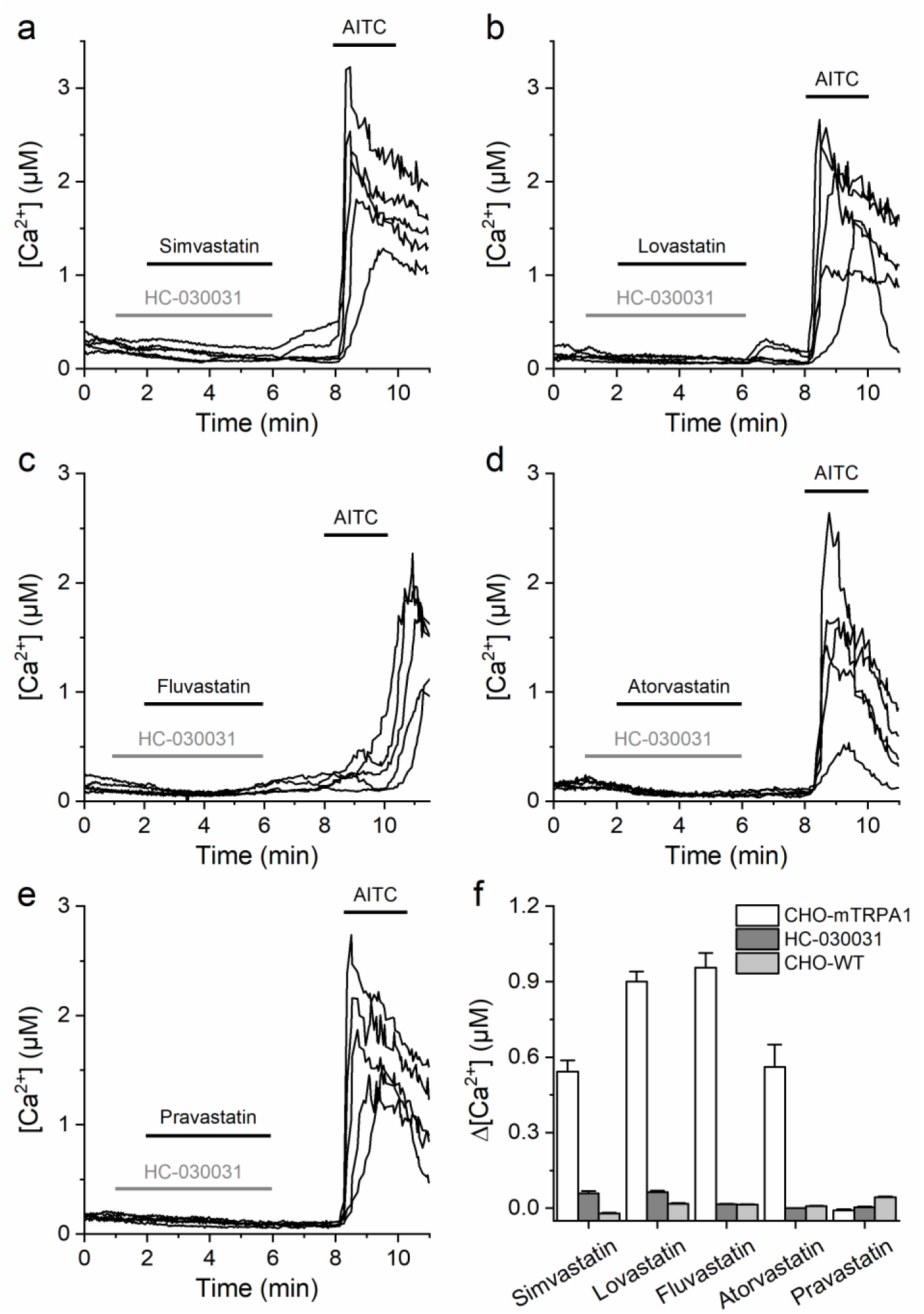
mTRPA1 responses to statins are inhibited by HC-030031. (a-e) Representative traces of [Ca^2+^] recorded in CHO-mTRPA1 cells showing the lack of activation for simvastatin (10 µM, a), lovastatin (20 µM, b), fluvastatin (200 µM, c), atorvastatin (400 µM, d), and pravastatin (800 µM, e) in the presence of TRPA1 inhibitor HC-030031 (100 µM). (f) Average [Ca^2+^] change induced by statins in CHO cells stably expressing mTRPA1 (white bar), in the presence of HC-030031 (dark gray bar), and in CHO cells not expressing TRPA1 (light gray bar). Statins were applied at concentrations of 10, 20, 200, 400, and 800 µM for simvastatin, lovastatin, fluvastatin, atorvastatin, and pravastatin, respectively. The data is represented as mean ± s.e.m. (n > 100).

**Figure 4 Supplement 1.**
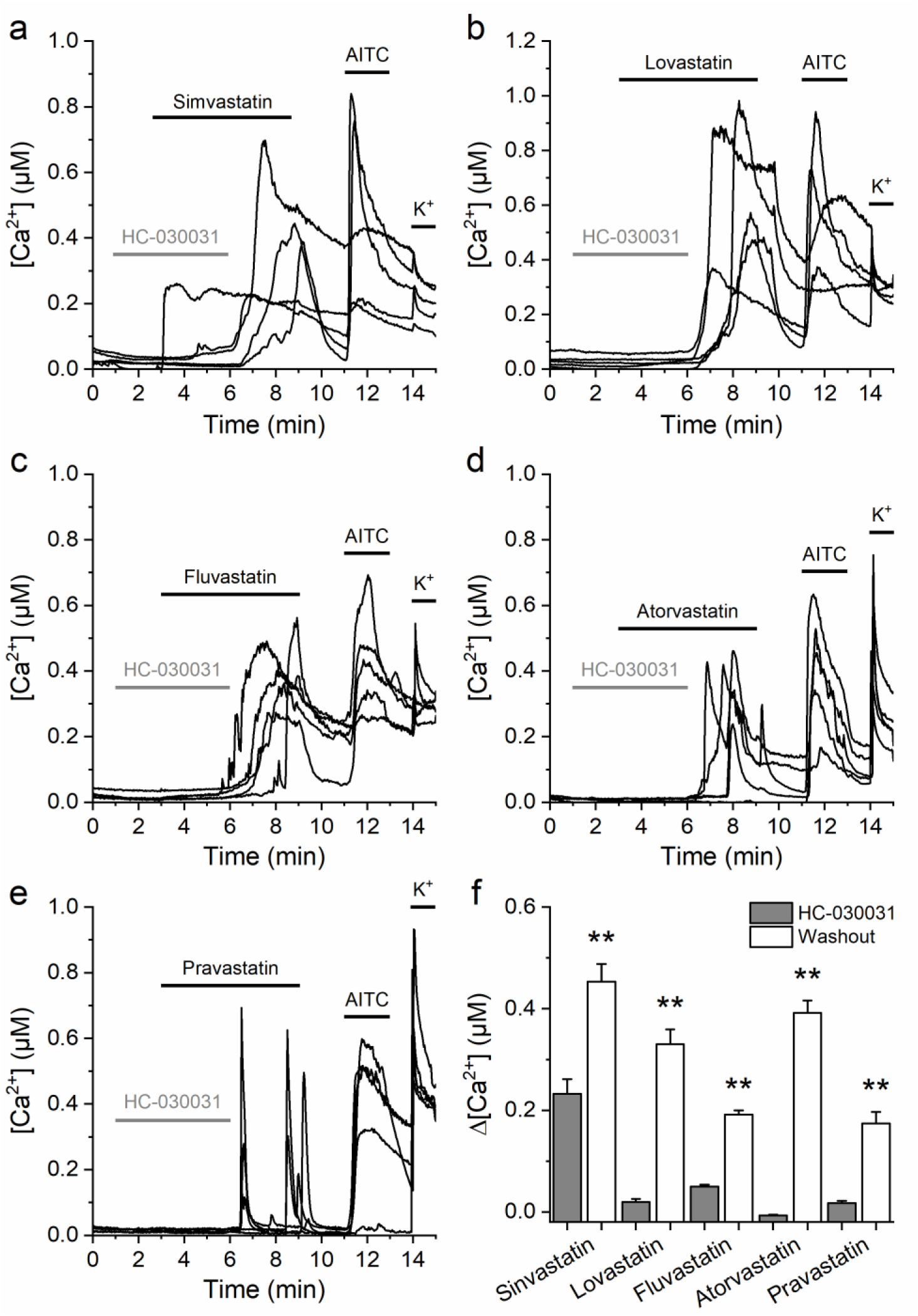
HC-030031 attenuates TRPA1 statins responses in mouse sensory neurons. (a-e) Representative traces of [Ca^2+^] responses of mouse DRG neurons to (a) 10 µM simvastatin; (b) 20 µM lovastatin; (c) 120 µM fluvastatin; (d) 360 µM atorvastatin; (e) 800 µM pravastatin and 30 µM AITC and high K^+^ (50 mM). (f) Average [Ca^2+^] amplitude change evoked by statins in the presence of HC-030031 (gray bar) and after removal of HC-030031 (washout; white bar). Data are shown as mean ± s.e.m for n = 127, 103, 224, 121, and 75 cells that were positive to simvastatin, lovastatin, fluvastatin, atorvastatin, and pravastatin, respectively, as well as to AITC and K^+^.

